# Symmetry structures in dynamic models of biochemical systems

**DOI:** 10.1101/2020.01.27.922005

**Authors:** Fredrik Ohlsson, Johannes Borgqvist, Marija Cvijovic

## Abstract

Symmetries provide a powerful concept for the development of mechanistic models by describing structures corresponding to the underlying dynamics of biological systems. In this paper, we consider symmetries of the non-linear Hill model describing enzymatic reaction kinetics, and derive a class of symmetry transformations for each order *n* of the model. We consider a minimal example consisting in the application of symmetry based methods to a model selection problem, where we are able to demonstrate superior performance compared to ordinary residual-based model selection. Finally, we discuss the role of symmetries in systematic model building in systems biology.

## 1 Introduction

The development of mathematical models is crucial in data-driven fields where the mechanism of the underlying system is of interest. In systems biology, mechanistic models of ordinary differential equations (ODEs) are often constructed to describe the change in abundance of an intracellular component such as mRNA or proteins over time. A proposed biological mechanism is typically combined with the law of mass action [1], yielding polynomial models. Under certain assumptions, e.g. regarding the relative abundance of different components, the models can be simplified giving rise to other types of non-linear rate equations which are common in enzyme kinetics [2]. A classic example is Michaelis–Menten kinetics or, more generally, the Hill equation describing the dynamics of a reaction forming a product, catalysed by an enzyme, in a situation where the concentration of the substrate is substantially higher than that of the enzyme [1]. The rate equations are the building blocks in the construction of mechanistic models in systems biology where each model implicitly proposes an underlying mechanism for the system at hand.

The prevailing strategies for constructing mechanistic models are based on data using a *top down*-approach. Given an experimental time series describing the change in the quantity of an intracellular component over time, numerous methods for model selection are based on *residual analysis* [3] using the *least squares* [1] cost function measuring the Euclidean distance between the measured data and the model predictions. Several proposed models are then evaluated and the one that minimises the cost function is selected. Other more sophisticated methods include the Akaike Information Criteria [3, 4, 5], Bayesian model selection [3, 4, 5], cross validation [6, 7] and bootstrapping methods [3, 8, 5] which for example take the model complexity in terms of the number of parameters into account. All these *statistical* methods rely on data (implying that experimental design is an integral part of model selection [9, 10]) which limits their applicability in cases when data is scarce or when several models describe the data equally well in terms of the residual analysis e.g. due to experimental errors which are large compared to the intrinsic variation across candidate models.

Model development can alternatively be conducted using mathematical analysis without any experimental data in a *bottom up*-approach. This is traditionally used e.g. in population dynamics [11], where the methodology consists in comparing different mathematical models of the same system [12] in terms of their agreement with properties derived from prior knowledge of the system rather than statistical measures. The most common tool for analysing ODE models in biology is *linear stability analysis* [1, 13] where the long term behaviour of the model is determined by linearising the system around its steady states. However, this asymptotic behaviour is often insufficient for completely determining the structure of the underlying system. An alternative technique for analysing a system of ODEs is to consider the set of symmetries of its solutions [14, 15]. The mathematical framework for such methods is that of group theory and representation theory, and more generally differential geometry. Symmetry methods have been used to classify ODE models according to their symmetry groups [16] and, conversely, identifying symmetries realised in a system allows for a constructive approach to modelling where the symmetries are made manifest in constructing the mechanistic model.

The symmetry framework is well-established and enormously successful for model construction in mathematical physics (e.g. as the foundational principle of the standard model of elementary particle physics [17, 18]). In fact, it has also found applications within mathematical biology such as animal locomotion [19], but is virtually unused in reaction kinetics modelling. Since the framework incorporates intrinsic properties of a system at all time scales, a symmetry based methodology could arguably represent an untapped potential for systems biology in the context of model development in particular and the analysis of dynamical models in general.

The aim of this paper is to elucidate the role of symmetries in systems biology by demonstrating a minimal example of the application of symmetry methods to model selection in enzyme kinetics. Specifically, provided a time series of the concentration of a substrate of an enzyme over time, and a number of candidate kinetic models describing the data approximately equally well in terms of the residuals, we apply a symmetry based method to select the model that is best able to represent the time series data. As the methodology is not commonly used in systems biology, we begin by establishing the framework and deriving a certain class of symmetries of the Hill models used in enzyme kinetics. Subsequently, we evaluate the proposed method applied to model selection among a set of three candidate Hill models. Finally, we discuss the benefits, validity and limitations of the proposed methodology.

## 2 Theoretical background

### 2.1 The Hill model

The Hill class of models, describing the enzymatically catalysed conversion of a substrate to a product, are defined by the ODE

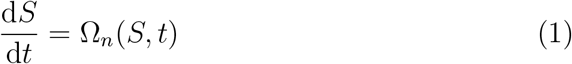

with

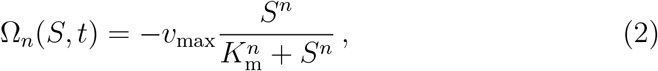

where *n* ∈ ℕ_+_ is the order of the model, *S* is the substrate concentration and *t* is the time. The parameters *v*_max_ and *K*_m_ correspond respectively to the maximum reaction rate and the substrate concentration at half of the maximum reaction rate. In all cases, physical solutions to (1) satisfy *S ≥* 0 ensuring that Ω_*n*_(*S, t*) is well defined.

Symmetry properties of the models are most easily analysed in terms of dimensionless time and concentration

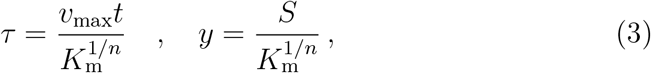

in terms of which the model (1) becomes

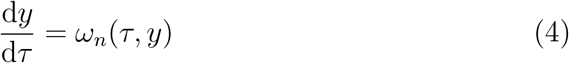

with

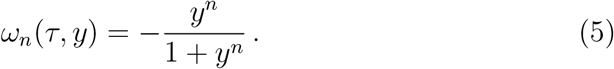

### 2.2 Symmetry transformations

A point transformation with parameter *є* ∈ ℝ acting on the (*τ, y*)-plane,

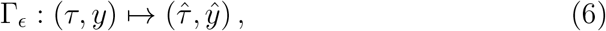

is a symmetry of the Hill model if it maps a solution of (4) to another solution. In other words the set of solutions of (4) is closed under the action of a symmetry Γ_*ϵ*_. The family of such symmetry transformations parameterised by *ϵ* forms a (representation of a) one-parameter Lie group.

There is no time dependence in the expression (5) for the derivative of the concentration, implying that time translation is a manifest symmetry of the theory. The corresponding point transformations are

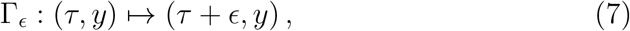

under which *ω_n_* are invariant for all model orders *n*.

In the context of elucidating structural properties of a model *ω_n_* from its symmetries, we will also consider point transformations Γ_*ϵ*_ which form a representation of a one-parameter Lie group but which are not symmetries of the model in the sense that the set of solutions is not closed under Γ_*ϵ*_.

It can be shown that the point transformation

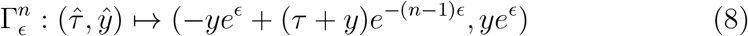

is a symmetry of the Hill model *ω_m_*(*τ, y*) of order *m* for *m* = *n* but not for *m* ≠ *n*. The symmetry transformation (8) is therefore unique to the Hill model of order *n*, which means that it can be used to distinguish between different Hill models. The action of the symmetry transformation on solutions to the Hill models of order *n* = 1, 2, 3 is illustrated in Fig. 1.

**Figure 1.**
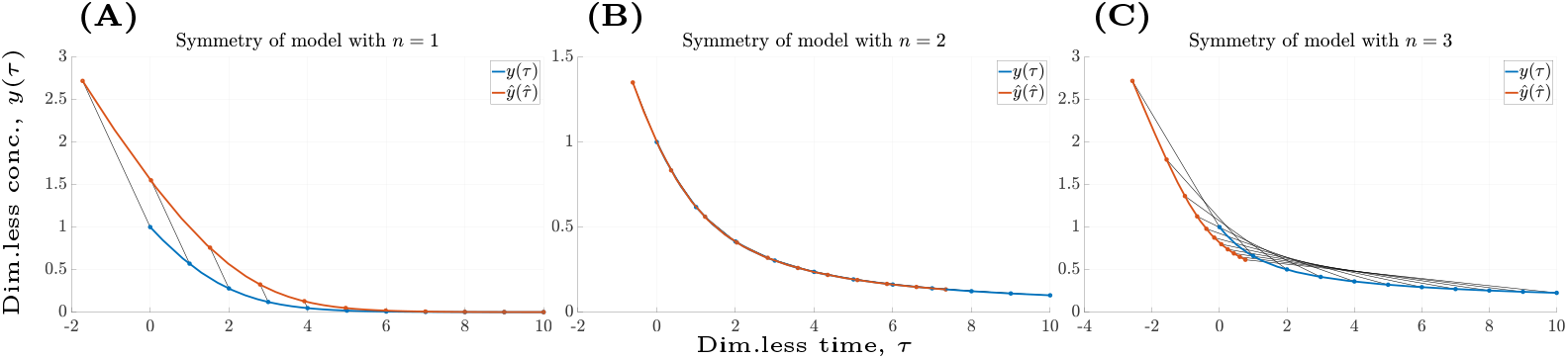
Action of symmetries. The action of the transformation 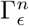 in (8) on solutions to the model *ω_n_*(*τ, y*) for **(A)** *n* = 1, **(B)** *n* = 2 and **(C)** *n* = 3. The action maps a solution *y*(*τ*) (blue) to a different solution 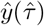 (red) for *n* = 1 and *n* = 3. For *n* = 2 the solution is invariant under the action of 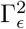 corresponding to symmetry which acts trivially on the space of solutions.

## 3 Symmetry based model selection

### 3.1 Method description

Given an experimentally acquired time series and a set of candidate models, the model selection problem consists in determining which candidate model is best able to describe the experimental data. In situations where several models fit the data approximately equally well in the least-square sense, they may still be differentiated by the extent to which they capture the global structure of the time series.

One way to achieve a comparison of this structural agreement is by using the fact that the space of solutions to a model is closed under the action of a symmetry transformation of that model, but not under generic transformations in (*τ, y*)-space. Consequently, the true model generating the time series should have the property that the least square-error is (approximately) invariant if the following steps are conducted. Initially, we apply a symmetry transformation Γ_*ϵ*_ to the data, then a model is fitted to the transformed data, the inverse transform is applied to the model and finally the least-square residuals are computed for the original time series. (The invariance is exact in the in the limit of vanishing errors.)

Conversely, if a symmetry transformation Γ_*ϵ*_ of an incorrect candidate model is applied in the same way the transformation will distort the time series and the quality-of-fit is expected to decrease. In particular, for a one-parameter group of symmetries we expect that the residuals will increase as a function of the parameter *ϵ* (at least locally in a neighbourhood of *ϵ* = 0). The effect on the quality-of-fit resulting from the procedure described above is illustrated in Fig. 2.

**Figure 2.**
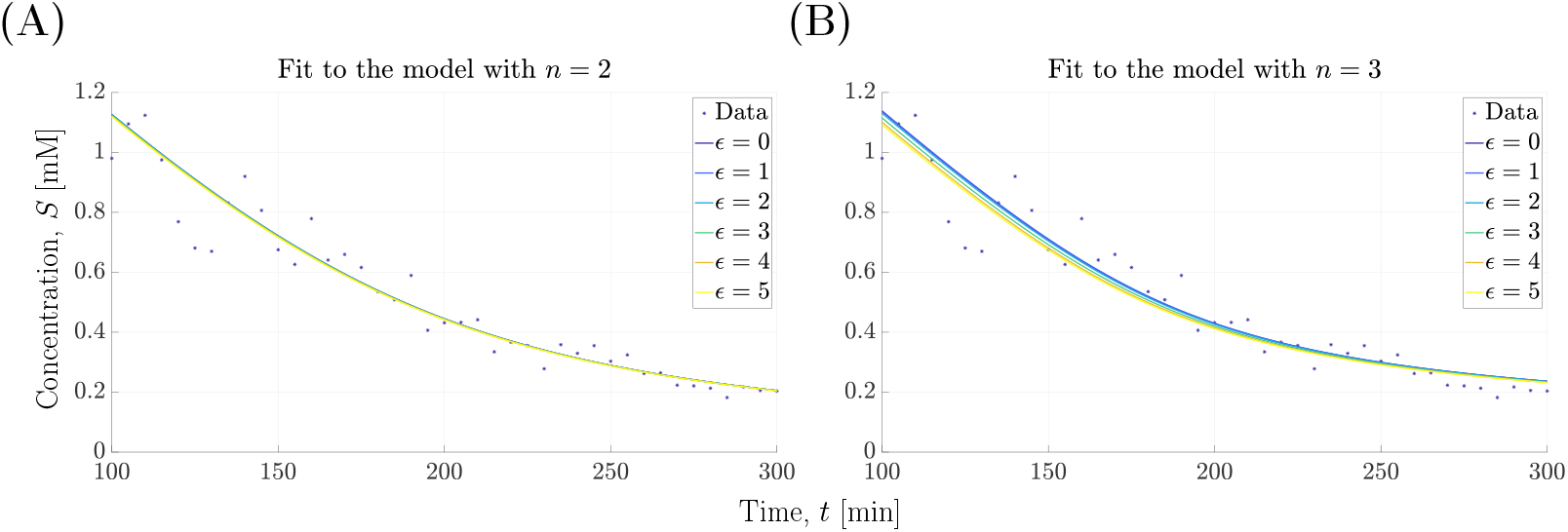
Quality-of-fit to transformed data. Hill models of order *n* = 2, 3 are fitted to data, simulated using a second order model, after application of the transformation 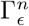. The inverse transform of the resulting fit (solid lines) is shown for increasing values of the transformation parameter *ϵ* for **(A)** *n* = 2 and **(B)** *n* = 3. The deterioration of the quality-of-fit for the model *n* = 3 results from 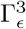 not being a symmetry of the underlying model generating the data.

The information about the dependence of the quality-of-fit on the transformation parameter *ϵ* can be used to complement the information obtained from the ordinary model fitting procedure. Thus, the purpose of the method for model selection described here is not to replace the common approach in systems biology, but rather to augment it using structural information about the candidate models (in the form of their symmetries) to provide additional information regarding their ability to represent a data set.

### 3.2 Evaluation setup

To evaluate the symmetry based model selection methodology, we consider a setup where a time series of substrate concentrations is simulated using a Hill model of order *n*_Sim_. Subsequently, a number of candidate Hill models, of different orders *n*_Fit_ are fitted to the simulated data using the “classic” least squares approach and the symmetry based methodology described above.

The classic approach is based on the *root mean square (RMS)*, *ρ*_0_ (41), where the selection criteria is that the model with the best fit, i.e. smallest value of *ρ*_0_, is selected. To calculate the statistical significance of the fitting of the candidate models to a single generated time series, the evaluation procedure is repeated *N* times and confidence intervals (CI) of the fits at the one standard error (SE) level are calculated. In this setting, models can be distinguished when their confidence intervals are not overlapping.

The symmetry based methodology is based on the RMS *ρ*(*ϵ*) (40) as a function of the transformation parameter *ϵ*. As in the the classic case, confidence intervals of the RMS-values are calculated as the evaluation is repeated *N* times. The selection criteria for the symmetry based methodology is that the model with the lowest RMS-value as *ϵ* increases is selected, significance requiring that the confidence intervals of the candidates do not overlap.

It should be noted that it is not obvious what range of the transformation parameter *ϵ* is required in order to differentiate between the candidate models using the symmetry based approach, or indeed if it is at all possible. In the examples considered in this paper, the range of *ϵ* is extended until the RMS curve *ρ*(*ϵ*) of each model reaches steady state. If no such state is obtained, the range is extended until convergence becomes prohibitively slow for the nonlinear optimiser or separation of candidate models is considered apparent.

## 4 The symmetry based methodology out-performs residual based fitting

The evaluation procedure described in the previous section is implemented for three cases *n*_Sim_ = 1, 2, 3. For each case, the candidate models *n*_Fit_ = 1, 2, 3 are fitted to a simulated time series using both the classic and symmetry based methods. The procedure is repeated *N* = 5 times and the corresponding confidence intervals are calculated. For the implemented noise levels and values of the kinetic parameters, i.e. *v*_max_ and *K*_m_, used in the simulations, the classic quality-of-fit is similar for all candidate models (Fig. 3).

**Figure 3.**
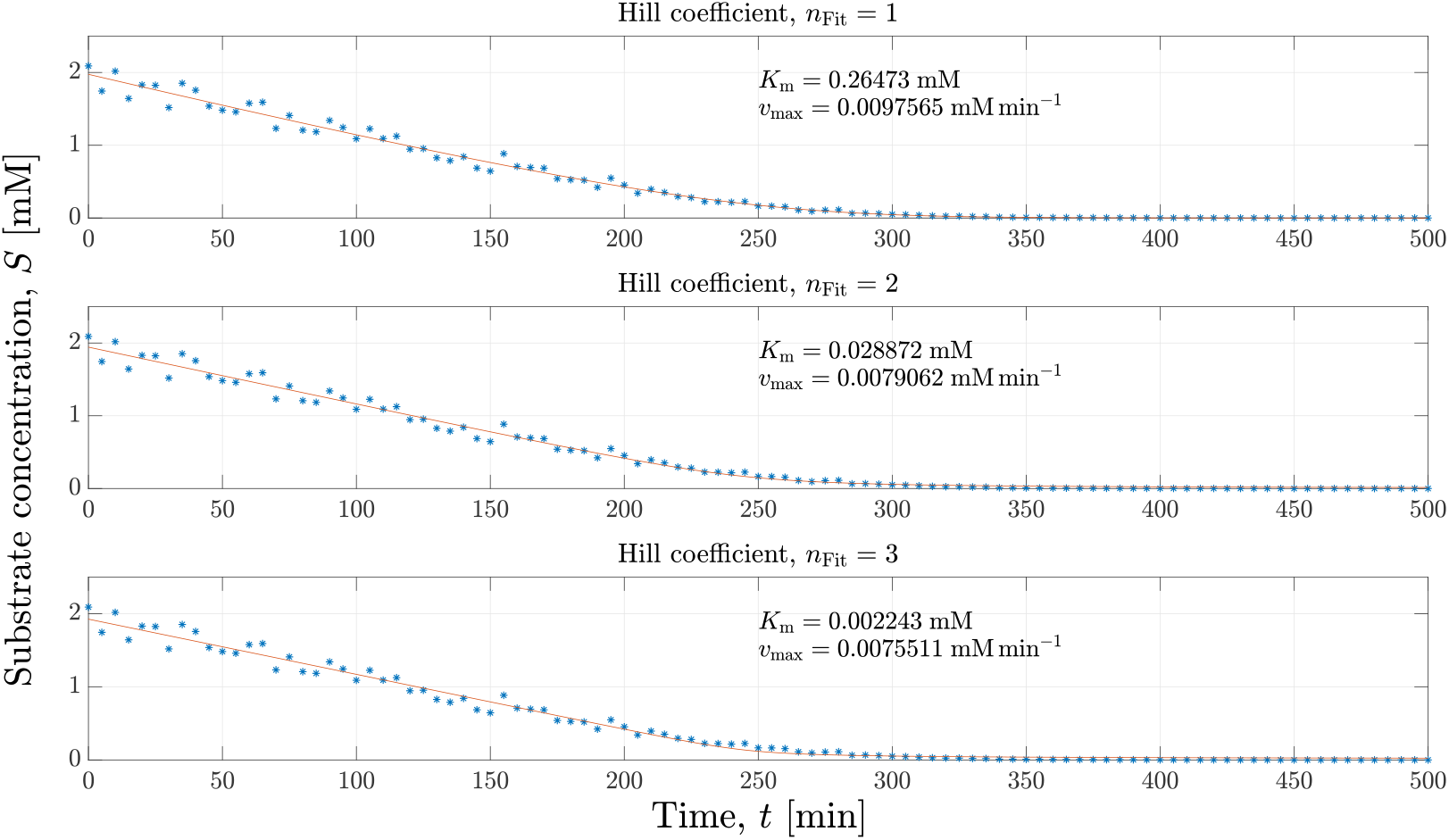
Individual fits of three Hill models. Three candidate models *n*_Fit_ = 1, 2, 3 (red) are fitted to the same simulated time series data with *n*_Sim_ = 1 (dashed blue) generated using a log-normal error-model with parameters *σ* = 0.1, *v*_max_ = 0.0102 mM min^*−*1^, *K*_m_ = 0.30 mM and *S*_0_ = 2 mM.

### 4.1 Data generated by the model with n_Sim_ = 1

In the *n*_Sim_ = 1 case it is possible to reject the *n*_Fit_ = 3 model but not to distinguish between the *n*_Fit_ = 1, 2 models using the classic approach, while the symmetry-based methodology selects the true model (Fig 4A-4B). The classic approach cannot distinguish between the *n*_Fit_ = 1, 2 models as the confidence intervals of the RMS *ρ*_0_ overlap (Fig 4A). However, the symmetry-based methodology clearly rejects the models with *n*_Fit_ = 2, 3 and selects the true model with *n*_Fit_ = 1 on the interval *ϵ ∈* [0, 5] (Fig 4B).

**Figure 4.**
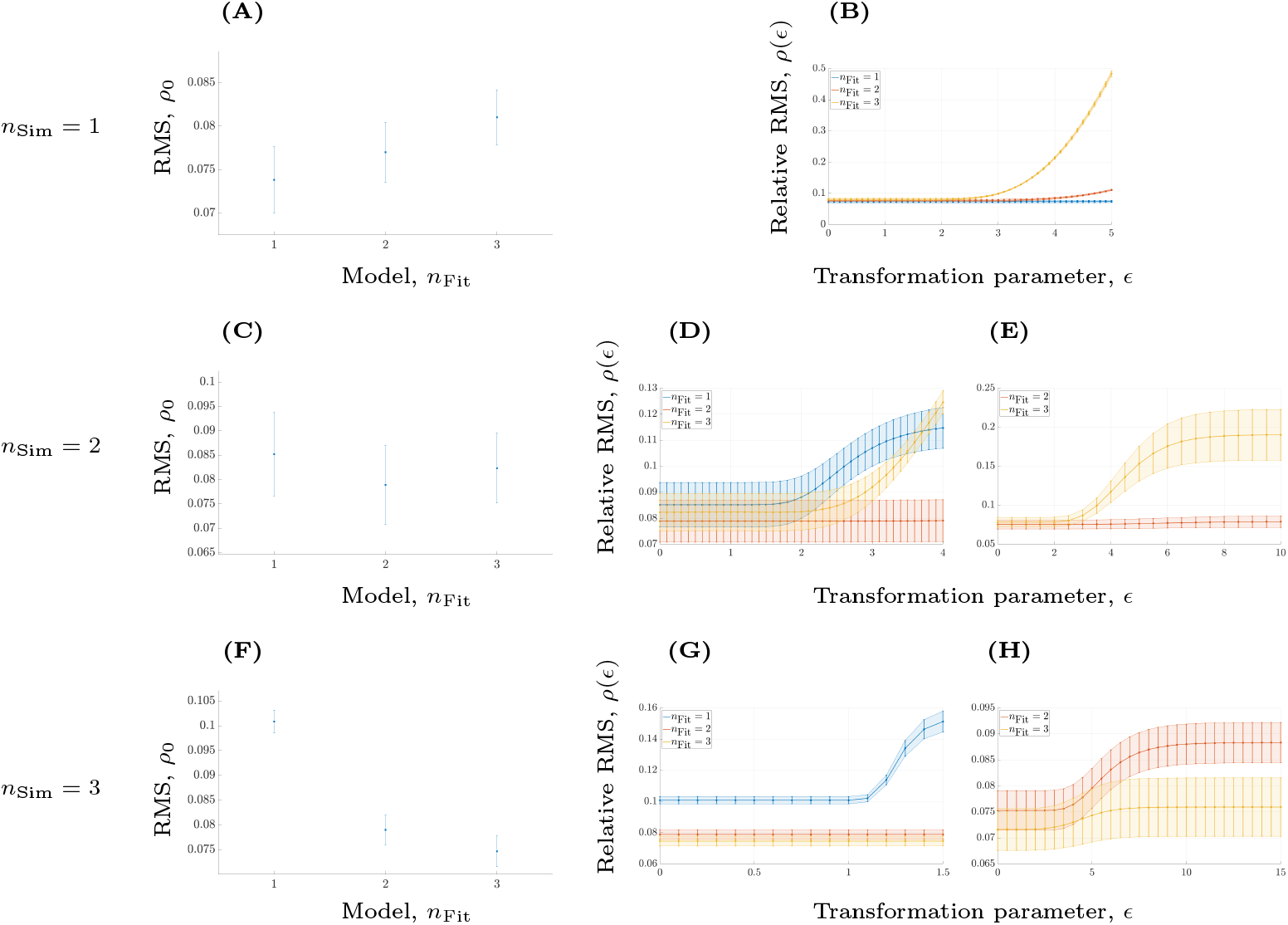
Model selection with distinct symmetries compared to the classic residual based approach. From the top to the bottom row, the data is generated with *n*_Sim_ = 1, 2, 3 using a log-normal error-model with parameters *σ* = 0.1, *v*_max_ = 0.0102 mM min^*−*1^, *K*_m_ = 0.30 mM and *S*_0_ = 2 mM. **(A)** The residual based RMS measure *ρ*_0_ fails to significantly distinguish between the *n*_Fit_ = 1, 2 models, but rejects the *n*_Fit_ = 3 model, for the data sets with *n*_Sim_ = 1. **(B)** Over the range *ϵ ∈* [0, 5] the symmetry based RMS measure *ρ*(*ϵ*) indicates that *n*_Fit_ = 1 is significantly better than *n*_Fit_ = 2, 3 for the data sets with *n*_Sim_ = 1. **(C)** The residual based RMS measure *ρ*_0_ fails to significantly distinguish between the *n*_Fit_ = 1, 2, 3 models for the data sets with *n*_Sim_ = 2. **(D)** Over the range *ϵ ∈* [0, 4] the symmetry based RMS measure *ρ*(*ϵ*) indicates that *n*_Fit_ = 2 is significantly better than *n*_Fit_ = 1, 3 for the data sets with *n*_Sim_ = 2. **(E)** Over the range *ϵ ∈* [0, 10] the symmetry based RMS measure *ρ*(*ϵ*) rejects the model with *n*_Fit_ = 3 and selects the model with *n*_Fit_ = 2 for the data sets with *n*_Sim_ = 2. **(F)** The residual based RMS measure *ρ*_0_ fails to significantly distinguish between the *n*_Fit_ = 2, 3 models but it can reject the first model with *n*_Fit_ = 1 for the data sets with *n*_Sim_ = 3. **(G)** Over the range *ϵ ∈* [0, 1.5] the symmetry based RMS measure *ρ*(*ϵ*) draws the same conclusion as the classic approach based on *ρ*_0_ in (F). In other words, the model with *n*_Fit_ = 1 is rejected while the methodology cannot distinguish between the *n*_Fit_ = 2, 3 models for the data sets with *n*_Sim_ = 3. **(H)** Over the range *ϵ ∈* [0, 15] the symmetry based RMS measure *ρ*(*ϵ*) rejects the model with *n*_Fit_ = 2 and selects the model with *n*_Fit_ = 3 for the data sets with *n*_Sim_ = 3.

### 4.2 Data generated by the model with n_Sim_ = 2

In the *n*_Sim_ = 2 case, the classic approach cannot distinguish between the models, as the confidence intervals of the RMS fitting overlap (Fig. 4C), while the symmetry based methodology selects the true model. Over the range *ϵ* ∈ [0, 4] the confidence intervals of the various models clearly separate using the symmetry based methodology (Fig. 4D) and the correct model with *n*_Fit_ = 2 is selected as it has the lowest RMS-value *ρ*(*ϵ*). In fact, this effect is exaggerated when the range of the transformation parameter is increased to *ϵ* ∈ [0, 10] (Fig. 4E) and it is evident in this case that the true model would be selected using the symmetry based approach.

### 4.3 Data generated by the model with n_Sim_ = 3

As in the previous cases, the symmetry based methodology outperforms the classic approach for *n*_Sim_ = 3. The classic approach rejects the first model with *n*_Fit_ = 1 while it cannot distinguish between the *n*_Fit_ = 2, 3 models as their confidence intervals overlap (Fig. 4F). For a short range of the transformation parameter *ϵ* ∈ [0, 1.5], the symmetry based methodology reaches the same conclusion (Fig. 4G). Thus, for small values of the transformation parameter *ϵ* the symmetry based methodology rejects the first model while it cannot distinguish between the other models as their confidence intervals of *ρ*(*ϵ*) overlap. However, by increasing the range of the transformation parameter to *ϵ* ∈ [0, 15] it is clear that the true model with *n*_Fit_ = 3 is selected and that the incorrect model with *n*_Fit_ = 2 is rejected (Fig. 4H).

## 5 Discussion

The main purpose of using symmetries in systems biology is the construction of mechanistic models. Fundamental properties of a given system can be described by their corresponding symmetries (e.g. energy conservation corresponds to invariance under time translations), and it is therefore of interest to be able to deduce the symmetries governing the system directly from the available data. By studying which symmetries a system obeys, it is possible to derive the corresponding dynamic models from those symmetries (e.g. in the form of the Lagrangian in analytical mechanics) [15, 20, 21].

By deriving symmetries of the Hill equation, a minimal example of the use of symmetries on the very building blocks of kinetic modelling in systems biology is presented. Moreover, we demonstrate that symmetries reveal intrinsic properties of a system of interest by presenting an example of a methodology for selecting Hill models based on a single time series. In fact, with one time series of substrate concentration over time no single model of the candidates corresponding to *n*_Fit_ = 1, 2, 3 can be identified using classic model fitting while the symmetry based methodology identifies the correct one in all cases (Fig. 4). We also validate the underlying assumption of the methodology, namely that the symmetries of the candidate models must be distinct by implementing the common translation symmetry (Fig. 7). Thus, this provides a minimal example of the fact that symmetries can be used to deduce intrinsic properties of a system where little data is available in a way regular model fitting cannot.

Importantly, the symmetry based model selection is not based on the assumption that any of the candidate models is in fact the correct model of the underlying system. If all evaluated transformations Γ_*ϵ*_ cause a significant increase in *ρ* with the transformation parameter *ϵ*, the conclusion is to reject all of the transformations at hand as symmetries of the system. Conversely, if several symmetries are identified any mathematical model of the system must respect all of them. The symmetry based evaluation described in the present paper should therefore more accurately be considered as the first step in a systematic model construction as oppose to a narrow methodology for selecting candidate models.

In cases where multiple time series are available it is possible to estimate the log-likelihood function and thereby use statistical methods such as the AIC or BIC criteria for model selection. However, if the actual underlying mechanisms of the studied system is of interest then symmetries can provide novel insights that classic model selection methodologies cannot. Therefore, symmetries are not meant to replace the already existing statistical methodologies for model selection but rather to complement them in the construction of mechanistic models.

A natural continuation of the work presented here is the automatisation of the methodology for identifying model symmetries in an algorithmic fashion, using numerical or computer algebra methods [15], to allow for systematic model structure identification for larger models describing the dynamics of e.g. large intracellular pathways. Such an automatisation requires the formulation of a criteria for the range of the transformation parameter *ϵ* in order to determine whether or not a certain transformation constitutes a symmetry. In addition, the methodology relies on Taylor expansions locally around *ϵ* ≈ 0 and it is not evident when the derived transformation ceases to be accurate. Furthermore, as discussed above, the range of the transformation parameter is crucial when using the symmetry based methodology as a means of selecting one model among multiple candidates. For example, over the range *ϵ* ∈ [0, 1.5] in the case of the data set generated with the model with *n*_Sim_ = 3, the methodology cannot distinguish between the models with *n*_Fit_ = 2 and *n*_Fit_ = 3 (Fig. 4G) while over the range *ϵ* ∈ [0, 15] (Fig 4H) the correct model corresponding to *n*_Fit_ = 3 is selected and the incorrect model with *n*_Fit_ = 2 is rejected.

As the ultimate goal of systems biology is to gain mechanistic understanding of complex cellular processes, symmetries constitute a forceful constituent in modelling where the underlying process is of interest. This study serves as an example of how this very potent methodology can be introduced into dynamic modelling in systems biology. As the symmetry framework is well-established in physics, the prospects of constructing, understanding and analysing models using symmetries in systems biology are exciting.

## Acknowledgements

This work was supported by Swedish Agency for Strategic Research (grant nr. IB13-0022) to MC. The authors gratefully acknowledge Philip Gerlee for valuable discussions of the work described and for suggesting improvements of the manuscript.

## A Supplementary material

### A.1 The Hill model

The Hill model describes the conversion of a substrate *S* into a product *P*. The reaction is catalysed by an enzyme *E* where the substrate binds to an active site of the enzyme forming a substrate-enzyme complex *C*. The complex then forms the product from the substrate in an irreversible reaction before dissociating from the product. In the general setting, it is possible to assume that an enzyme has *n* ∈ ℕ_+_ active sites corresponding to the assumption that *n* substrate units are required in order to form one unit of the product. Furthermore, under certain conditions it is possible to assume that it is the binding of the first substrate unit that is the *rate-limiting step*, implying that as soon as one the first substrate unit binds to the enzyme the other units bind immediately. These conditions are described from a kinetic point of view by the following set of reactions.

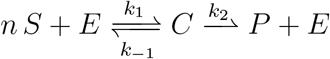

In the above reactions, *k*_1_ is the rate constant for the binding of the substrate units to the enzyme, *k_−_*_1_ is the rate constant for the dissociation of the substrate units from the enzyme and *k*_2_ is the rate constant describing the conversion of the substrate units to the product. Assuming that the *law of mass action* holds, the dynamics of these reactions is governed by

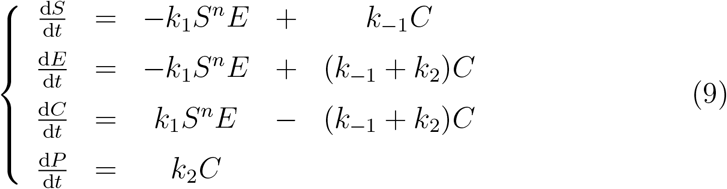

with initial conditions

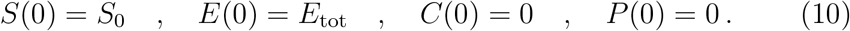

Note that *the total number of enzymes E*_tot_ = *ϵ* + *C* appears as the initial condition for the unbound form of the enzyme *E* since there is no complex initially and it is clear that *E*_tot_ is conserved

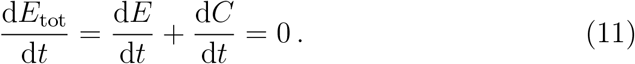

Assuming that the amount of enzyme is much smaller than the amount of substrate, i.e. *S*_0_ ≫ *E*_tot_, implying that the enzymes are always saturated, then it is possible to motivate the assumption

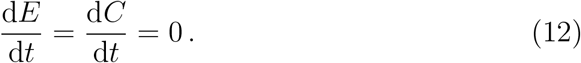

Substituting this equation into the system of equations above yields

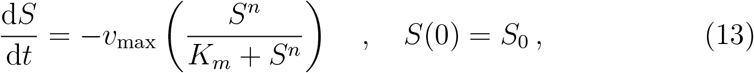

which is referred to as the *Hill equation*, in which we have introduced the constants

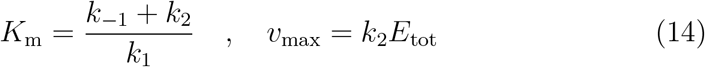

and the number of active sites *n* ∈ N_+_ is called the *Hill coefficient*, or the order of the Hill model.

The non-dimensionalisation of the Hill equation is obtained by introducing

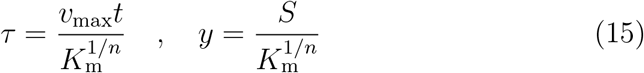

and substituting these two dimensionless components into the original model (13) yields the following dimensionless version of the equation describing the consumption of the substrate

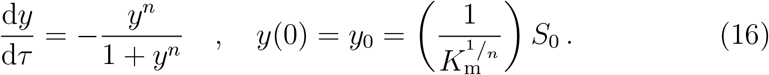

It is worth emphasising that the only parameter in the dimensionless model (16) is the initial condition *y*_0_.

### A.2 Simulation methodology for generating the data

In simulating data for the Hill model, we have used kinetic parameters for the enzyme *β*-lactamase I from the organism *Bacillus cereus* [22, 23]. The reported parameters corresponding to the full enzymatic system (9) are *k*_1_ = 0.068 mM^*−*1^min^*−*1^, *k_−_*_1_ = 0.0136 min^*−*1^ and *k*_2_ = 0.0068 min^*−*1^. In the simplified Hill model (13) this corresponds to a value of *K*_m_ = 0.30 mM and we have implemented a value of *E*_tot_ = 1.5 mM for the total enzyme concentration resulting in the maximal reaction rate *v*_max_ = 0.0102 mM min^*−*1^. All simulations use an initial substrate concentration of *S*_0_ = 2 mM and a log-normal error-model

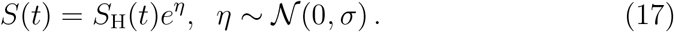

In the above equation, *S*(*t*) corresponds to the simulated data at time *t*, *S*_H_(*t*) corresponds to the underlying process at time *t*, given by the solution the ODE-model (13), and *η* is the error drawn from a normal distribution with standard deviation *σ*. In all simulations, we have implemented a noise level of 10%, i.e. *σ* = 0.1. An example of simulated data is shown in Fig. 5.

**Figure 5.**
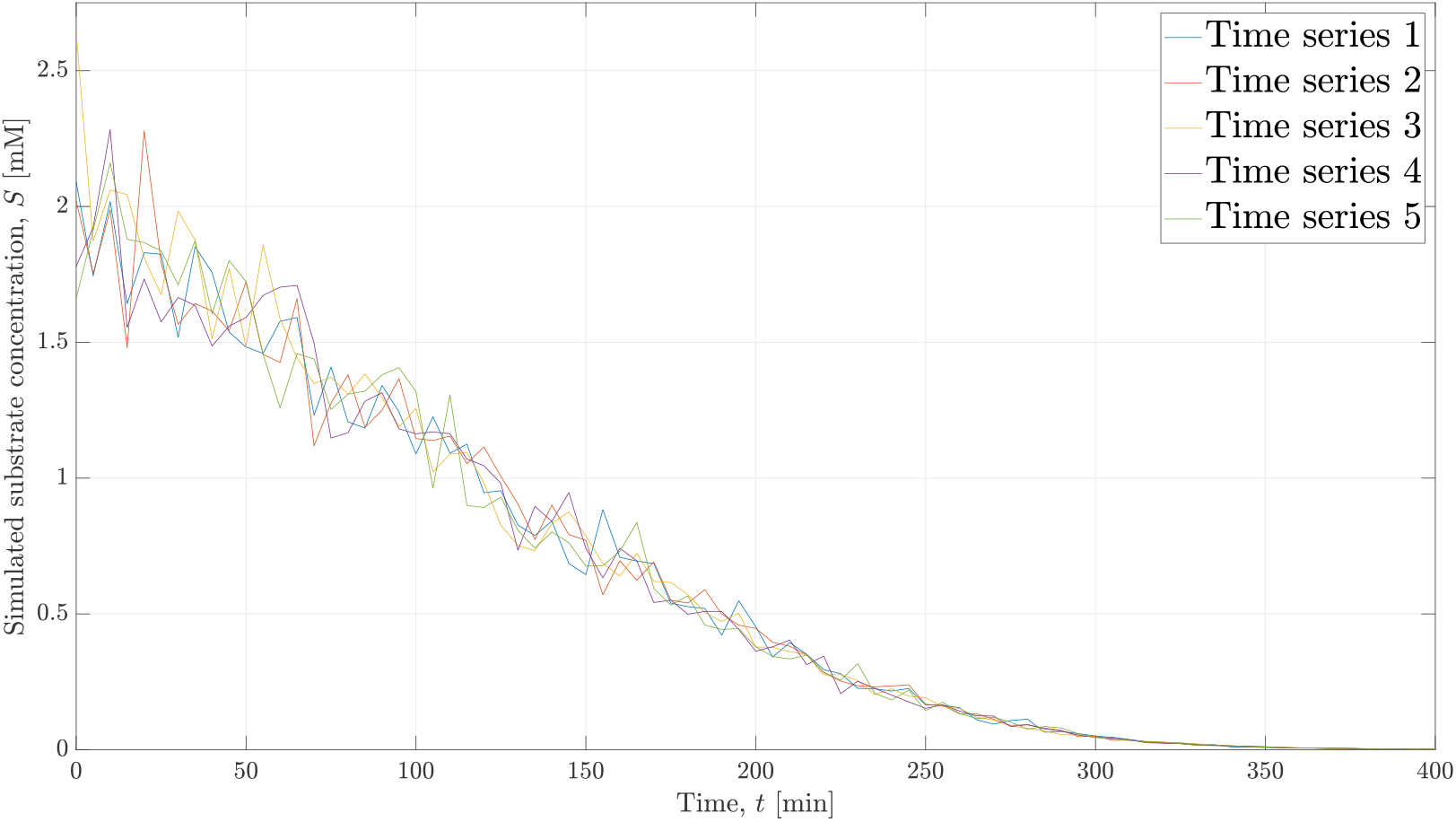
Simulated data. Five time series with substrate concentration over time are presented. The data is simulated with a log-normal error-model with parameters *σ* = 0.1, *v*_max_ = 0.0102 mM min^*−*1^, *K*_m_ = 0.30 mM and *S*_0_ = 2 mM.

The choice to implement a log-normal error-model (17), as opposed to a simpler additive error-model, requires some motivation. Firstly, for concentrations close to zero it is possible to obtain negative data-points with an additive model which is avoided with the log-normal model. Secondly, in applications it is often the case that errors associated with measurements for high concentrations are often higher than the corresponding measurement-errors for low concentrations [24]. This effect is captured in the log-normal error-model but not in the additive model, in which the absolute errors are of equal size across the entirety of the time series.

### A.3 Symmetries of first order ODE’s

In this section we summarise the general theory for Lie symmetries of a single first order ODE

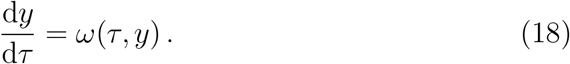

A solution of (18) is a curve *y*(*τ*) in the (*τ, y*) plane and a point transformation is a map Γ: ℝ^2^ → ℝ^2^ defined by its action on an arbitrary point

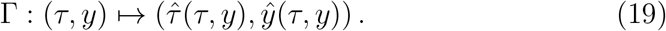

A point transformation constitutes a symmetry of the ODE (18) if it maps the set of solutions to itself, that is if

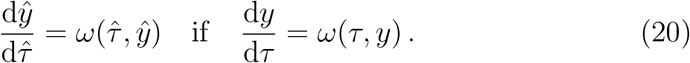

We consider exclusively sets of symmetry transformations,

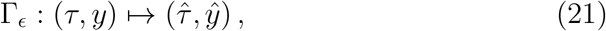

parameterised by a number *ϵ* ∈ ℝ, which are diffeomorphisms of ℝ^2^ and form a representation of a (local) one-parameter Lie group *G*. For such representations Γ_0_ is the trivial transformation, and there exists a neighbourhood *U* of *ϵ* = 0 such that Γ_*δ*_Γ_*ϵ*_ = Γ_*δ*+*ϵ*_ for *δ, ϵ* ∈ *U* and 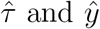 can be represented as Taylor series in *ϵ* in *U*. In particular, this implies that the inverse of a transformation is obtained as

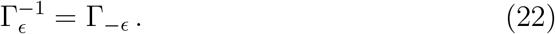

The set of points (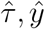) obtained by the action of the Lie group *G* on (*τ, y*) is called the orbit of (*τ, y*). At the point (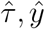) the vector tangent to the orbit is (*ξ*(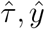), *η*(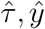)) where

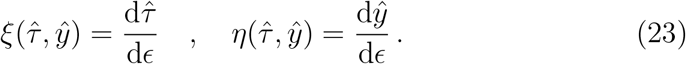

In particular, the existence of a Taylor series expansion around *ϵ* = 0 implies that

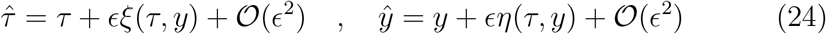

with

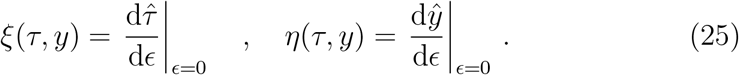

Since the symmetry transformations Γ_*ϵ*_ are diffeomorphisms of ℝ^2^, the assignment of (*ξ, η*) is smooth and defines a vector field on ℝ^2^ by

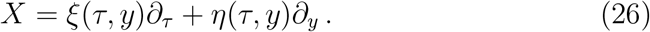

The vector field generates the symmetry transformation Γ_*ϵ*_ through

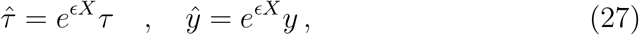

where *e*^*ϵX*^ is the equivariant exponential map satisfying

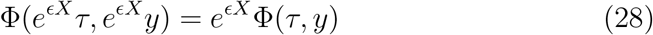

for an arbitrary function Φ: ℝ^2^ → ℝ. The vector field *X*, called the infinitesimal generator of Γ_*ϵ*_, contains all information required to reconstruct the corresponding transformations.

Consequently, the equation (18) admits a one-parameter Lie group of symmetries generated by (26) if *ξ*(*t, y*) and *η*(*t, y*) satisfy the linearised symmetry condition

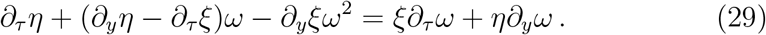

### A.4 Symmetries of the Hill model

The class of symmetries primarily considered in the present paper are obtained by using an Ansatz linear in both *τ* and *y* for the components *ξ*(*τ, y*) and *η*(*τ, y*) of the generating vector field *X*, according to

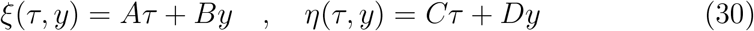

for the Hill model *ω_n_*(*τ, y*) in (5). The linearised symmetry condition (29) takes the form

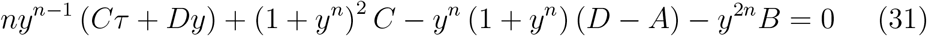

which for *n* ∈ ℕ_+_ has the general solution

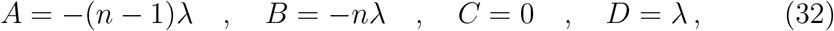

where *λ* ∈ ℝ is an arbitrary constant. Up to simultaneous rescalings, the components of the tangent vector are therefore

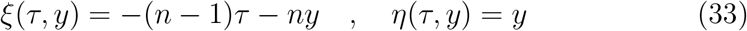

and the corresponding generating vector field is given by

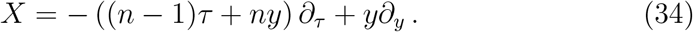

The coordinate transformation generated by (34) is obtained by the exponential map as

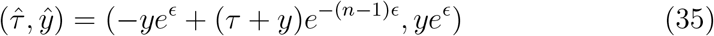

where *ϵ* is the transformation parameter.

In addition to the linear symmetries described above, which are different for models of different values of *n*, the Hill models (5) are all manifestly invariant under a time translation transformation

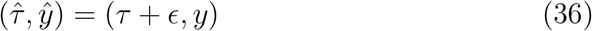

for all *n* ∈ ℕ_+_. The transformation is generated by the vector field

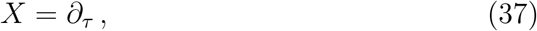

with components *ξ*(*τ, y*) = 1 and *η*(*τ, y*) = 0, which trivially satisfy the linearised symmetry condition since *∂_τ_ ω_n_* = 0. Since the Hill model is a first order ODE, its solution is uniquely determined by a choice of initial condition at (say) *τ* = 0. Consequently, any non-trivial symmetry transformation can be obtained as equivalent time translation with a suitable choice of transformation parameter.

### A.5 Model selection method

The purpose of the symmetry based method for model selection is to incorporate global structural properties of the time series in the comparison of candidate models. Symmetry transformations preserve the space of solutions to the ODE model, whereas general coordinate transformations do not. Consequently, if we apply a symmetry transformation Γ_*ϵ*_ of the candidate model *ω*(*τ, y*) to the (non-dimensionalised) time series data, perform a least square fit to the transformed data to obtain a solution of the candidate model, and then apply the inverse transformation Γ_*−ϵ*_ to the fitted model, the result will be a (different) solution to the model *ω*(*τ, y*). If the transformation Γ_*ϵ*_ is also a symmetry of the model generating the time series data, the residuals between the original time series data and the resulting solution should be approximately independent of *ϵ*. In the limit of vanishing errors in the time series data the independence becomes exact.

Conversely, if a transformation Γ_*ϵ*_ which is not a symmetry of the true model is applied in the same way, it will distort the transformed time series data, causing a reduction in the quality-of-fit of the candidate model. The dependence of the residuals on the transformation parameter *ϵ* is non-linear, but in a neighbourhood of *ϵ* = 0 we expect the residuals of the fit to be increasing since *ϵ* = 0 corresponds to the trivial transformation which introduces no distortion. Throughout this paper, we use the ordinary RMS error, *ρ*(*ϵ*) (40), for the residuals to quantify the quality-of-fit.

Since the symmetries of each candidate Hill model are most conveniently implemented in the dimensionless coordinates (*τ, y*) according to (15), a preliminary step in the model selection process outlined above is to estimate the values of *K*_m_ and *v*_max_ from the simulated data 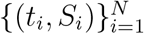, using ordinary nonlinear least-square optimisation, and compute the non-dimensionalised data 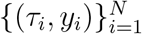.

In order to avoid introducing a dependence on the non-dimensionalisation, which differs depending on the candidate model, *ρ*(*ϵ*) is always computed in the original dimensional context. Furthermore, to obtain a meaningful comparison between the effects of symmetry transformations of different candidate models we also normalise the scale of the parameter *ϵ* so that *ϵ* = 1 corresponds to the initial data point (*τ*_1_*, y*_1_) being shifted by (at least) 50% of the time series range in both the *τ* and the *y* direction,

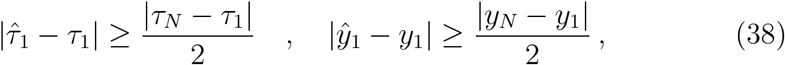

where (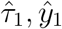) is the result of applying 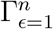 to (*τ*_1_*, y*_1_).

Using the fact that the solution to any first order ODE is uniquely determined by a choice of initial condition, we introduce the notation

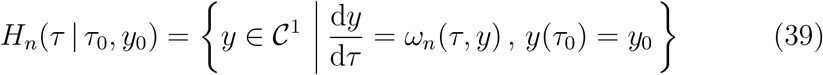

for a solution to the Hill model of order *n* ∈ ℕ_+_. Given the time series 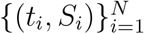 of substrate concentrations, a set of integers *n* defining the candidate Hill models and a corresponding set of representations 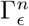 of one-parameter symmetry groups unique to each candidate model, the method for symmetry based Hill model selection can be described as follows. For each value *n* of the candidate model order:

1. Etimate parameters *K*_m_ and *v*_max_ from 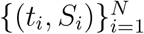
2. Compute non-dimensionalised time series 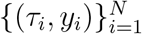
3. Normalise transformation parameter *ϵ*
4. For each value *ϵ* of the transformation parameter:
  i. Apply transformation 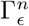 to the data

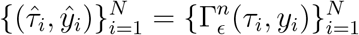
  ii. Set 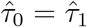 and determine least-square fit 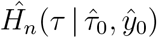 as

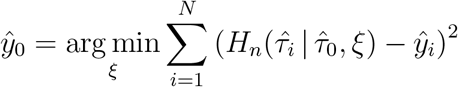
  iii. Apply the inverse transform 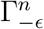 to the model

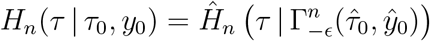
  iv. Evaluate the fit *ρ*(*ϵ*) of *H*_*n*_ to 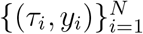

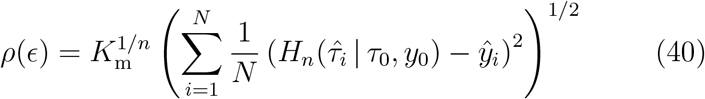

The different parts of Step 4 of the method are illustrated in Fig. 6.

**Figure 6.**
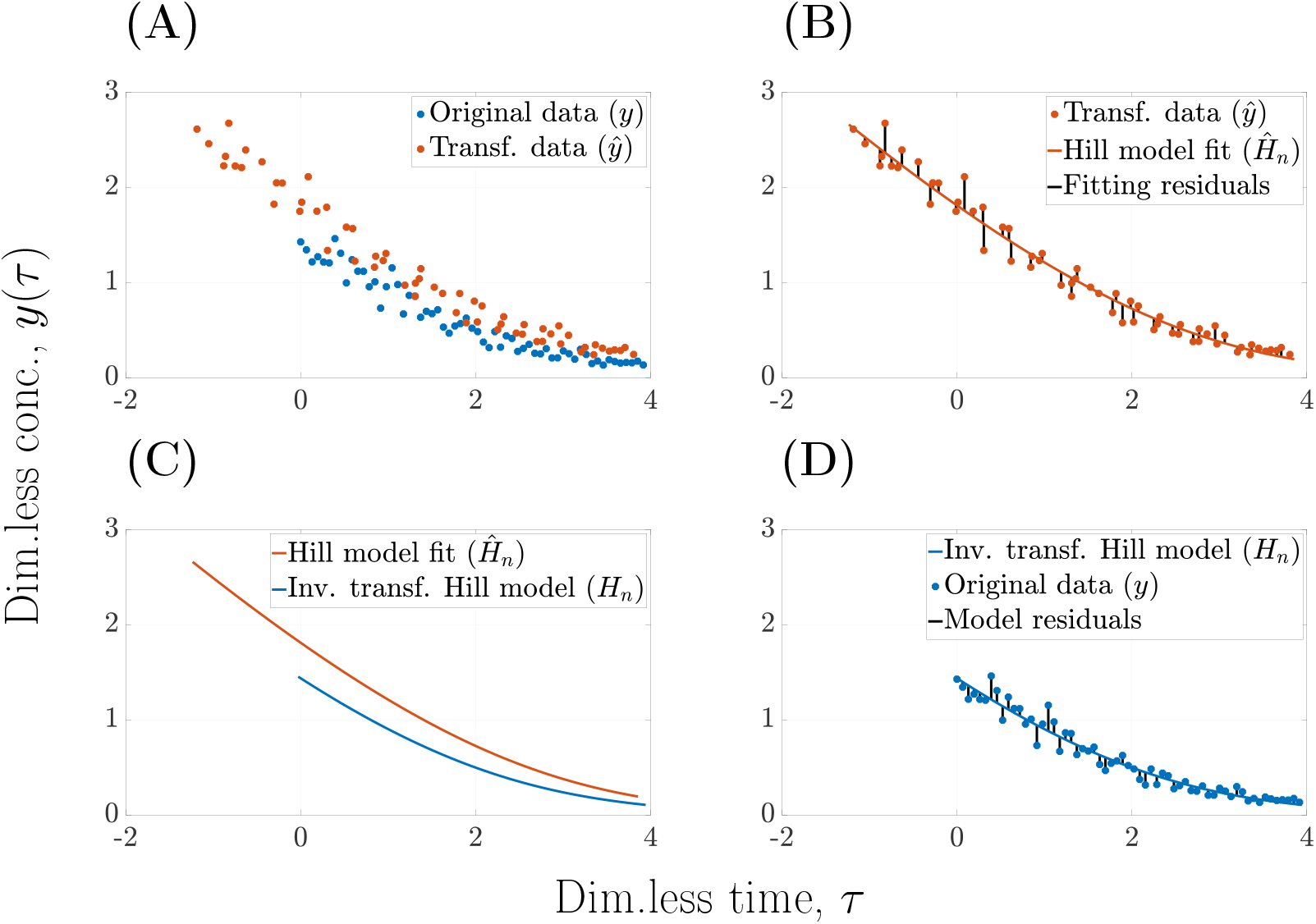
Illustration of the symmetry based method for model selection. The simulated dimensionless concentration *y*(*τ*) plotted versus dimensionless time *τ* for the *n* = 1 model. The point transformation implemented is the symmetry 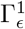 of the model. **(A)** The original (*y*) and the transformed (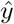) time series. **(B)** The transformed time series (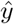), the fitted Hill model (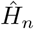) and the corresponding residuals. **(C)** The fitted Hill model (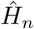) and its inverse transform (*H*_*n*_). **(D)** The inverse of the fitted Hill model (*H*_*n*_), the original data (*y*) and the corresponding residuals.

Using the function *ρ*(*ϵ*) we can express the ordinary RMS error of the classic approach as

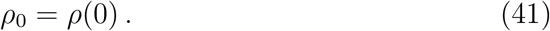

### A.6 Validation using the translation symmetry

In order to establish the validity of the symmetry based methodology we investigate the case of a point transformation which is not a symmetry for one of the candidate model but not the others. The time translation transformation Γ_*ϵ*_ in (7) is a symmetry of all Hill models, and it is therefore expected that the goodness-of-fit is approximately independent of the parameter *ϵ* for all model orders. We define the *relative RMS*, ∆(*ϵ*) as

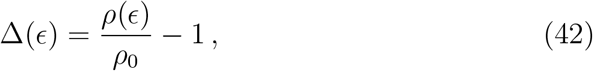

where the value ∆(*ϵ*) = 0 corresponds to the transformation having no effect on the fitting procedure, and perform the symmetry based fitting procedure.

As expected, the common translation symmetry does not distinguish between the candidate models. For all three data sets generated with the models *n*_Sim_ = 1, 2, 3 the confidence intervals of the relative RMS is of the order 10^*−*12^ centered around ∆ = 0 (Fig. 7). Numerical tolerance of the optimiser is set to 10^*−*12^ which suggests that ∆(*ϵ*) is zero to within numerical errors. Accordingly, the translation transformation (7) is confirmed as a symmetry of all models *n*_Fit_ = 1, 2, 3 incapable of distinguishing between the candidate models. This result validates the fundamental assumption of the symmetry-based methodology, that the symmetries Γ_*ϵ*_ of the candidate models must be distinct in order to differentiate between them. Furthermore, it provides a consistency check of the method using a symmetry transformation different from the specific transformations 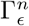 in (8) used to generate the results in Fig. 4.

**Figure 7.**
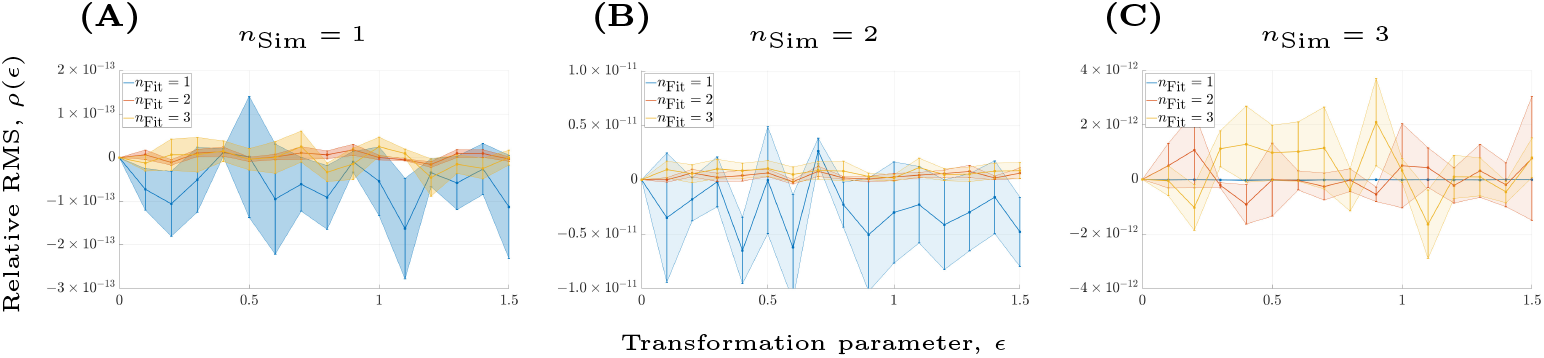
Symmetry based approach for the translation symmetry. In all three cases, the models with *n*_Fit_ = 1, 2, 3 are fitted to the simulated data over the range *ϵ* ∈ [0, 1.5]. The relative RMS ∆(*ϵ*) is plotted against the transformation parameter *ϵ* where the model selection is conducted with the common translation symmetry Γ_*ϵ*_: (*t, y*) ↦ (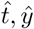) = (*t* + *ϵ, y*). The data is generated with the models corresponding to **(A)** *n*_Sim_ = 1, **(B)** *n*_Sim_ = 2 and **(C)** *n*_Sim_ = 3 respectively. The data is generated using a log-normal error-model with parameters: *σ* = 0.1, *v*_max_ = 0.0102 mM min^*−*1^, *K*_m_ = 0.30 mM and *S*_0_ = 2 mM. In all cases, the methodology cannot distinguish between any of the models as the relative RMS is within the range of the numerical tolerance, i.e *|*∆(*ϵ*)| ≈ 10^*−*12^. This result confirms the fact that Γ_*ϵ*_ is a symmetry of all three models.

